# STAPLER: Efficient learning of TCR-peptide specificity prediction from full-length TCR-peptide data

**DOI:** 10.1101/2023.04.25.538237

**Authors:** Bjørn P. Y. Kwee, Marius Messemaker, Eric Marcus, Giacomo Oliveira, Wouter Scheper, Catherine J. Wu, Jonas Teuwen, Ton N. Schumacher

**Author notes:** These authors contributed equally: Bjørn Kwee, Marius Messemaker.

## Abstract

The prediction of peptide-MHC (pMHC) recognition by αβ T-cell receptors (TCRs) remains a major biomedical challenge. Here, we develop STAPLER (Shared TCR And Peptide Language bidirectional Encoder Representations from transformers), a transformer language model that uses a joint TCRαβ- peptide input to allow the learning of patterns within and between TCRαβ and peptide sequences that encode recognition. First, we demonstrate how data leakage during negative data generation can confound performance estimates of neural network-based models in predicting TCR – pMHC specificity. We then demonstrate that, because of its pre-training and fine-tuning masked language modeling tasks, STAPLER outperforms both neural network-based and distance-based ML models in predicting the recognition of known antigens in an independent dataset, in particular for antigens for which little related data is available. Based on this ability to efficiently learn from limited labeled TCR-peptide data, STAPLER is well-suited to utilize growing TCR – pMHC datasets to achieve accurate prediction of TCR – pMHC specificity.

## Introduction

The recognition of peptide-MHC complexes by αβ T cell receptors (TCRs) forms the critical step in the initiation of a T cell-mediated adaptive immune response^1^. T cells acquire both the α and β chain of the TCR through somatic recombination of V(D)J gene segments and random non-templated nucleotide additions and deletions at the gene junctional boundaries, thereby forming the highly diverse CDR3α and CDR3β regions that make the most extensive physical contacts with the peptide antigen^2^ (**Fig. 1a**). Each of the many unique TCRs that can be generated through this process (estimated to be at least ∼3·10^11^)^3^ has the potential to recognize a distinct set of peptide-MHC complexes, allowing an individual’s collective T cell pool to cover an overwhelming diversity of such pMHC complexes^3–5^. This ‘distributed recognition capacity’ enables the joint T cell pool to recognize and clear pathogen infected cells that present foreign peptide antigens^6^, act as a modifier of the growth of tumors that present cancer (neo-)antigens^7^, and inadvertently start a self-destructive response against healthy cells that present self-peptides in the context of autoimmunity^8^.

**Fig. 1:**
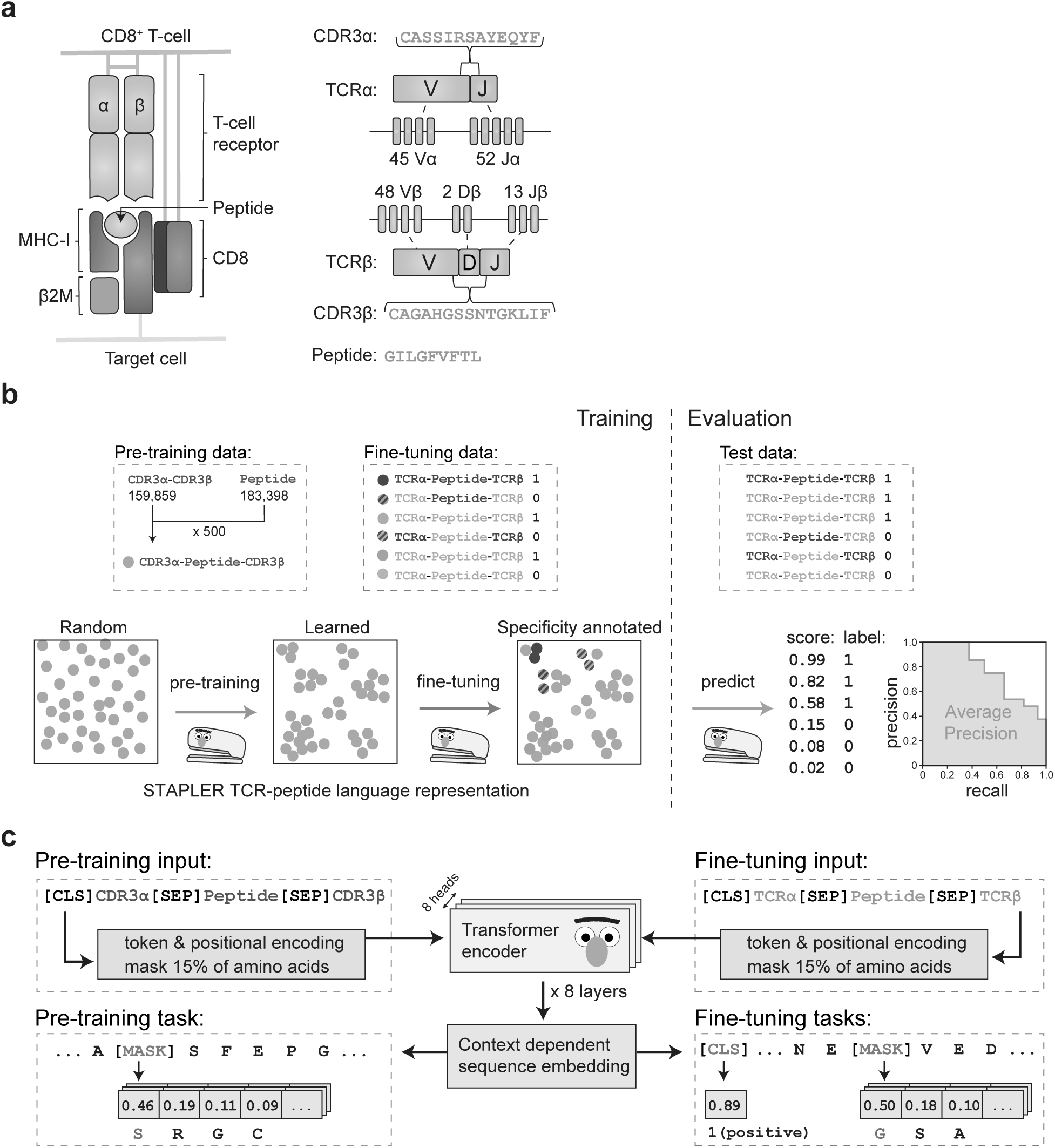
Shared TCR And Peptide Language bidirectional Encoder Representations from transformers (STAPLER) model. **a.** Schematic representation of the interaction between a CD8^+^ T cell-expressed αβ TCR and a target cell expressed peptide-MHC-I (pMHC) complex (left) and the process of TCR diversification through V(D)J recombination (right). Random non-templated nucleotide additions and deletions at the gene junctional boundaries form the highly diverse CDR3α and CDR3β regions that make the most extensive physical contact with peptides^2, 3^. **b**. STAPLER training (left) and STAPLER evaluation (right) pipeline. STAPLER was trained in two stages. In a first “pre-training” phase, large datasets of known MHC-I-presented peptides and CDR3 regions of TCRαβ pairs with unknown pMHC reactivity were used to learn language representations of peptide antigen and TCRαβ CDR3 amino acid sequences. In a second “fine-tuning” phase, these TCRαβ-peptide language representations were used to learn the prediction of TCR – pMHC specificity from a finite pool of labeled (i.e., with known reactivity or known lack of reactivity) TCRαβ-peptide pairs. STAPLER was evaluated using Average Precision (AP; area under the precision-recall curve) on a hold-out test dataset of labeled TCRαβ-peptide pairs. **c**. STAPLER is a Medium sized (8 heads, 8 layers) encoder from transformers model^16^ that was pre-trained using an MLM task (left) and fine-tuned using both a recognition label prediction task (indicated by the classification token ([CLS]) that reflects the reactivity score) and an MLM task (right). Pre-training was performed on random combinations of CDR3αβ pairs and peptides, and fine-tuning on both positive and negative full-length TCRαβ- peptide combinations. Reactive full-length αβ TCR-peptide pairs are labeled positive, non-reactive full-length αβ TCR-peptide pairs are labeled negative.

Because of the essential role of the TCR in a variety of human disease processes, efforts that aim to diagnose or modulate adaptive T cell responses would greatly benefit from the ability to predict TCR – pMHC reactivity from just their amino acid sequences. In such efforts, two discrete use-cases can be distinguished that are related to the class of peptide antigen involved. Specifically, a first class of use cases involves the prediction of the presence of T cell reactivity against previously identified T cell antigens (i.e., known antigens that can be included in model training). Next to that, a second class of use cases involve the prediction of T cell reactivity against previously unknown antigens. The latter use-case will, for instance, allow the prediction of T cell reactivity against the private neoantigens that arise as a consequence of somatic mutations in cancer tissue^7^, or T cell reactivity against pathogens not previously present in the human population^9^. In contrast, the former use case will allow the diagnosis of T cell reactivity against existing human pathogens, autoimmune disease-associated antigens, and shared cancer (neo-)antigens. In such efforts, the ability to call T cell reactivity may, for instance, be utilized to determine disease status (e.g., virus-positive or -negative), or to monitor the (activation) state of the pathogen-, autoimmune or tumor-reactive T cell compartment of interest.

Many of the prior models to infer pMHC specificity of TCRs have been trained to predict reactivity against a fixed set of predetermined peptide antigen categories (Supplementary Table **1**). However, as argued previously^10–15^, prediction of TCR reactivity against both known and unknown antigens will require a model that is based on a joint TCRαβ and peptide input (**Extended Data Fig. 1a**). The development of models that utilize such a joint TCRαβ-peptide input has to date been limited by the availability of labeled (i.e., known reactive or known non-reactive) TCRαβ-peptide data (**Extended Data Fig. 1c**). We hypothesized that, for two reasons, transformer language models^16^ that can be trained using the masked language modelling (MLM) task^17^, in which the identity of randomly masked amino acids is predicted from their surrounding sequence, would be particularly suited to address this challenge. Firstly, as the MLM task is self-supervised, it may be used to pre-train models to learn protein language representations from a (potentially large pool of) unlabeled protein sequences. In the case of TCR – pMHC prediction, this would involve the use of the large datasets of known MHC-presented peptides and the large datasets of TCRαβ pairs with an unknown pMHC reactivity to learn the structure of peptide antigens and of TCRαβ sequences. The resulting language representations could then allow the more efficient (i.e., requiring less data to reach the same predictive performance) learning of recognition patterns in the finite pool of TCRαβ-pMHC combinations with known reactivity (hereafter referred to as labeled TCRαβ-peptide data; **Fig. 1b, 1c**). This two-step approach is conceptually similar to the initial pre-training of models to learn general natural language representations from massive corpora of texts^17^, to subsequently fine-tune these pre-trained models for the efficient learning of supervised natural language tasks (e.g., predicting the class to which a specific text belongs)^17^. Secondly, next to the potential value of MLM-based pre-training, the masking of amino acids during model fine-tuning may force models to consider amino acid positions beyond the minimal amino acid positions that are required to predict the recognition label in the labeled fine tuning data. For example, a unique ‘code’ of a few amino acids in an individual peptide antigen may already suffice as a shortcut^18^ to predict the recognition label in the labeled fine-tuning data, thereby ignoring other residues that contribute to the interaction. By randomly masking peptide amino acids of such ‘codes’ during fine-tuning, MLM may thus force the model to develop a more holistic view of the peptide antigen. By the same token, the masking of TCR amino acids could allow a model to associate TCR sequence language with pMHC reactivity in the broadest possible sense. Based on the potential value of MLM to learn TCR – pMHC language, we set out to develop a transformer language model that uses a joint TCRαβ-peptide input to predict TCR-peptide specificity.

## Results

### Development and validation of STAPLER

In this model, called Shared TCR And Peptide Language bidirectional Encoder Representations from transformers (STAPLER), language representations from CDR3αβ and peptide sequences were first learned using an MLM pre-training task, exploiting a large pool of unlabeled CDR3αβ sequences and peptide sequences that we aggregated (pre-training performed using 79,929,500 random CDR3αβ - peptide combinations from 159,859 CD8^+^ T cell-derived CDR3αβ sequences and 183,398 9-mer peptides that bind MHC-I, **Fig. 1c** and **Extended Data Fig. 1b**). Next, the parameters of this pre-trained STAPLER model were fine-tuned on 23,544 labeled pairs of CD8^+^ T cell-derived full-length αβ TCRs and 9-mer peptides that bind MHC-I (**Extended Data Fig. 1c**), using both prediction of the TCR-peptide recognition label (i.e., a positive label for reactive pairs and a negative label for non-reactive pairs) and an MLM fine-tuning task (**Fig. 1c**). Importantly, to force learning of recognition patterns between TCR and peptide sequences, rather than recognition class (i.e., positive or negative label) occurrence bias of individual TCR sequences and peptide sequences^15, 19^, an issue that has been reported to result in data leakage in other fields^18, 20^, we ensured that each unique TCR and each unique peptide sequence occurred with equal frequency with either a positive or negative label in these data (see Methods, **Extended Data Fig. 3a**).

To validate the performance of STAPLER, a series of 5-fold cross-validation experiments was performed in which we benchmarked STAPLER against ERGO-AE-II_vj, ERGO-LSTM_vj, and NetTCR- 2.0^11, 12^—the three previously established models that use a joint TCR αβ-peptide input but that do not make use of a transformer language model. In addition, the performance of these different neural network models was compared to that of a simple feedforward neural network (ffwd) model, to evaluate the benefit of more complex neural network architectures. Model performance was analyzed using average precision (AP), a metric that estimates the fraction of predicted positives that will validate in wet-lab efforts across varying thresholds (**Fig. 1b**). Of note, this metric is insensitive to the class imbalance^21^ between positive (i.e., reactive) and negative TCR-peptide pairs that is common in typical biological samples^4^. Importantly, STAPLER outperformed the ERGO-AE-II_vj, ERGO-LSTM_vj, and NetTCR-2.0 models by +12.3% mean average precision (mAP), surpassing the improvement that the ERGO-II and NetTCR-2.0 models provide over the simple ffwd model (**Fig. 2a**).

**Fig. 2:**
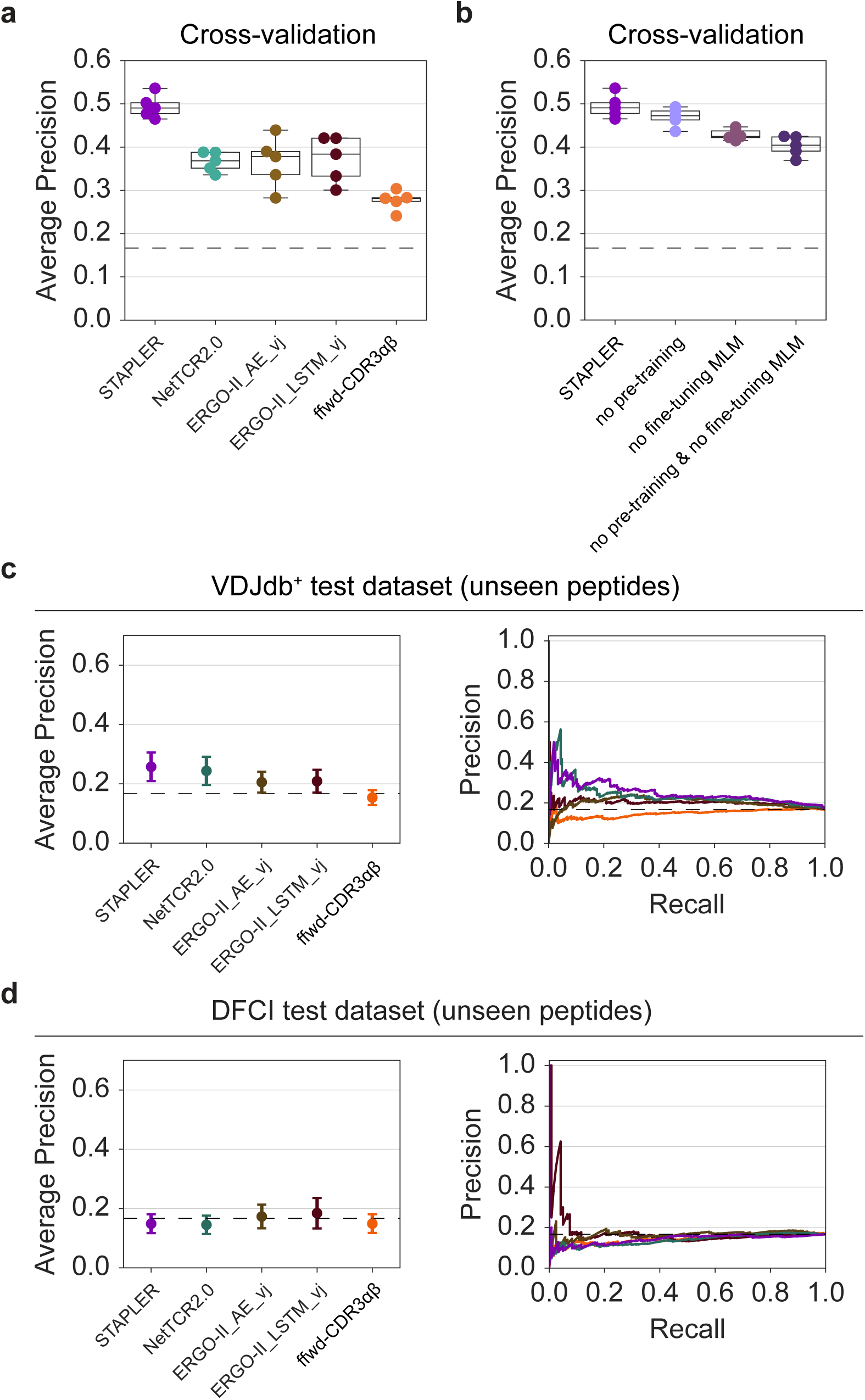
Cross-validation of STAPLER on the fine-tuning dataset and importance of negative data generation strategies. **a**. Comparison of STAPLER, NetTCR-2.0, ERGO-AE_vj, and ERGO-LSTM_vj in 5-fold cross-validation AP on the fine-tuning dataset. A simple ffwd model (CDR3-ffwd) was included as a baseline comparator. **b**. Average precision (AP) of STAPLER and indicated model variants in 5-fold cross-validation on the fine-tuning dataset. Note that pre-training and, in particular, fine-tuning MLM contribute to model performance. **a-b**. Dots show AP of each fold, boxes indicate model AP median and 25th/75th percentiles, whiskers show min/max except for outliers determined using the inter quartile range (IQR). Dashed line shows random performance (AP of 0.1667). **c-d**. mean average precision (mAP) (left), and precision across recall thresholds (right), on predicting recognition of unseen peptides antigens by unseen TCRs in the VDJdb^+^ test dataset (206 positive and 1,030 negative pairs, 16 unique unseen peptide sequences) (**c**) and of unseen peptides antigens by unseen TCRs in the DFCI external test dataset (121 positive and 605 negative pairs, 12 unique unseen peptides) (**d**). Dots show mAP and error bars show mAP 95% confidence intervals. Dashed line shows random performance (0.1667 AP).

Having established that STAPLER outperforms previously established methods in cross-validation, we set out to test the basis for this improved performance using defined model perturbations. First, to control for a possible bias in the fine-tuning TCR and peptide sequence data that could allow the model to predict the recognition label of TCR sequences or peptide sequences independent from the peptide or TCR that these sequences were paired with, we generated mock-up variants of STAPLER in which fine-tuning was performed while either omitting TCR information (thereby testing for bias in peptide sequences), or omitting peptide information (thereby testing for bias in TCRαβ sequences). While an intact STAPLER model (without pre-training, see Methods) showed an AP of 47% in cross-validation, AP dropped to random in the absence of either TCR or peptide information during model fine-tuning (**Extended Data Fig. 2a**), indicating that the performance of STAPLER is based on information that is obtained from paired TCR – pMHC sequence data. Furthermore, performance was optimal when using full-length αβ TCR sequences as input, as compared to either CDR3αβ only-, TCRα only-, TCRβ only-, or a V and J gene only-based models (**Extended Data Fig. 2b-d**), thereby supporting the notion that full-length αβ TCR input is preferable for TCR specificity prediction (**Extended Data Fig. 1a**).

To directly test to what extent the MLM pre-training and MLM fine-tuning tasks, the two main components introduced in STAPLER as compared to prior joint TCRαβ-peptide models (Supplementary Table **1**)^11, 12^, contributed to superior model performance, we assessed the performance of STAPLER upon the individual or joint ablation of these components. Importantly, ablation of the individual components reduced model performance numerically (pre-training) and significantly (MLM fine tuning task), and the combined ablation of pre-training and the MLM fine-tuning task resulted in a significant drop in model performance (**Fig. 2b**).

### Data leakage as a confounder in performance evaluation of TCR – pMHC specificity models

As discussed above, the two main use cases of TCR – pMHC specificity models are the prediction of TCR recognition of unknown epitopes, such as patient-specific antigens, and the predicted recognition of known epitopes, such as shared antigens. In view of the large size of the potential TCR – pMHC space and the currently extremely sparse experimental sampling of this space, it seems intuitive to focus current model use on the prediction of T cell reactivity against known antigens that can be included in model training. While for the majority of models, above-random performance on unknown peptides has indeed not been observed, such above-random performance on unknown peptides has been reported for the ERGO-II models^12^. To explore this aspect, and to understand whether dataset composition could influence such analyses, we evaluated the performance of STAPLER and the prior joint TCRαβ-peptide models using two test datasets, VDJbd^+^ and DFCI^22^. The VDJdb^+^ dataset is a collection of only positive TCR-peptide pairs (206 pairs) that were identified in different studies, in large part through MHC multimer staining, and this dataset both contains peptides that were present (i.e., seen) or absent (i.e., unseen) during model training (see Methods for definition of unseen peptide antigens and unseen TCR sequences). In contrast, the DFCI dataset is composed of both positive (121 pairs) and negative (605 pairs) TCR-peptide pairs that we identified in melanoma samples, and only contains peptides (melanoma antigens and viral antigens) that were unseen during model training.

While the DFCI dataset contains its own, experimentally measured, negative TCR-peptide pairs, such negative TCR-peptide pairs need to be artificially generated for datasets such as VDJdb^+^. Notably, most joint TCRαβ-peptide models have previously been evaluated on test datasets that contain negative TCR-peptide pairs that were generated by ‘mispairing’ peptides of each positive datapoint with other TCRs present in the dataset^12–15, 23^. However, it may be proposed that such mispairing with internal TCR sequences could result in data leakage. Specifically, such mispairing causes a higher proportion of the artificially generated negative pairs to contain TCRs that resemble TCRs present in the fine-tuning dataset than what would typically be present in a biological sample (data leakage class L3.2 in Ref. ^20^). Thus, in case models have the ability to detect that TCRs in such artificial negative pairs are likely to be reactive with their original cognate known antigen, rather than the peptide antigen that they have been mispaired with (**Extended Data Fig. 3b**), this would be predicted to result in an overestimate of model performance.

To explore whether performance estimates would be influenced by the type of negative dataset used, we first compared STAPLER, NetTCR-2.0, and the ERGO-II models with respect to their ability to predict TCR recognition of unknown antigens by unseen TCRs in the VDJdb^+^ and DFCI datasets. Notably, while all models showed a significantly (STAPLER and NetTCR-2.0) or numerically (the ERGO-II models) better-than-random performance on predicting the recognition of unseen peptides by unseen TCRs in the VDJdb^+^ dataset (**Fig. 2c**), such above-random performance was not observed for the DFCI dataset (**Fig. 2d**). To directly test whether the type of negative TCR-peptide pairs present in a dataset could explain the difference in model performance between the two datasets, peptides of each positive datapoint in VDJdb^+^ were mispaired with TCRs sampled from the measured negative pairs in the DFCI dataset. Vice versa, peptides of each positive datapoint in the DFCI dataset were mispaired with TCRs sampled from VDJdb^+^ (**Extended Data Fig. 3b**). Importantly, the better-than-random performance of all models on VDJdb^+^ largely disappeared when the DFCI TCRs were used for mispairing. The other way around, the performance of all models on the DFCI dataset increased when VDJdb^+^ TCRs were used for negative pair generation. Similarly, performance of all models on unknown antigens in the VDJdb^+^ dataset was also reduced when a second set of external TCRs (identified in oral cavity squamous cell carcinomas^24^) was used for negative data generation (VDJdb^+^-ETN, **Extended Data Fig. 3c-e**; also observed when using AUC as a metric **Extended Data Fig. 3f-h**). Together, these data demonstrate how data leakage in a commonly used strategy for negative data generation in TCR-pMHC prediction affects performance estimates, and describe a ‘best practice’ for the creation of test datasets for the evaluation of model performance.

### STAPLER outperforms other models in predicting the recognition of known antigens

To subsequently compare the different neural network models with respect to their ability to correctly identify novel TCRs that recognize known antigens, we evaluated performance using the VDJdb^+^-ETN dataset that avoids data leakage due to negative data generation. In this analysis, both the ERGO-II models and NetTCR-2.0 showed a numerical but non-significant increase in performance as compared to the ffwd baseline model. Notably, STAPLER outperformed all the prior models, with an improvement in AP of +9.6% mAP relative to ERGO-II_LSTM_vj (**Fig. 3a**).

**Fig. 3:**
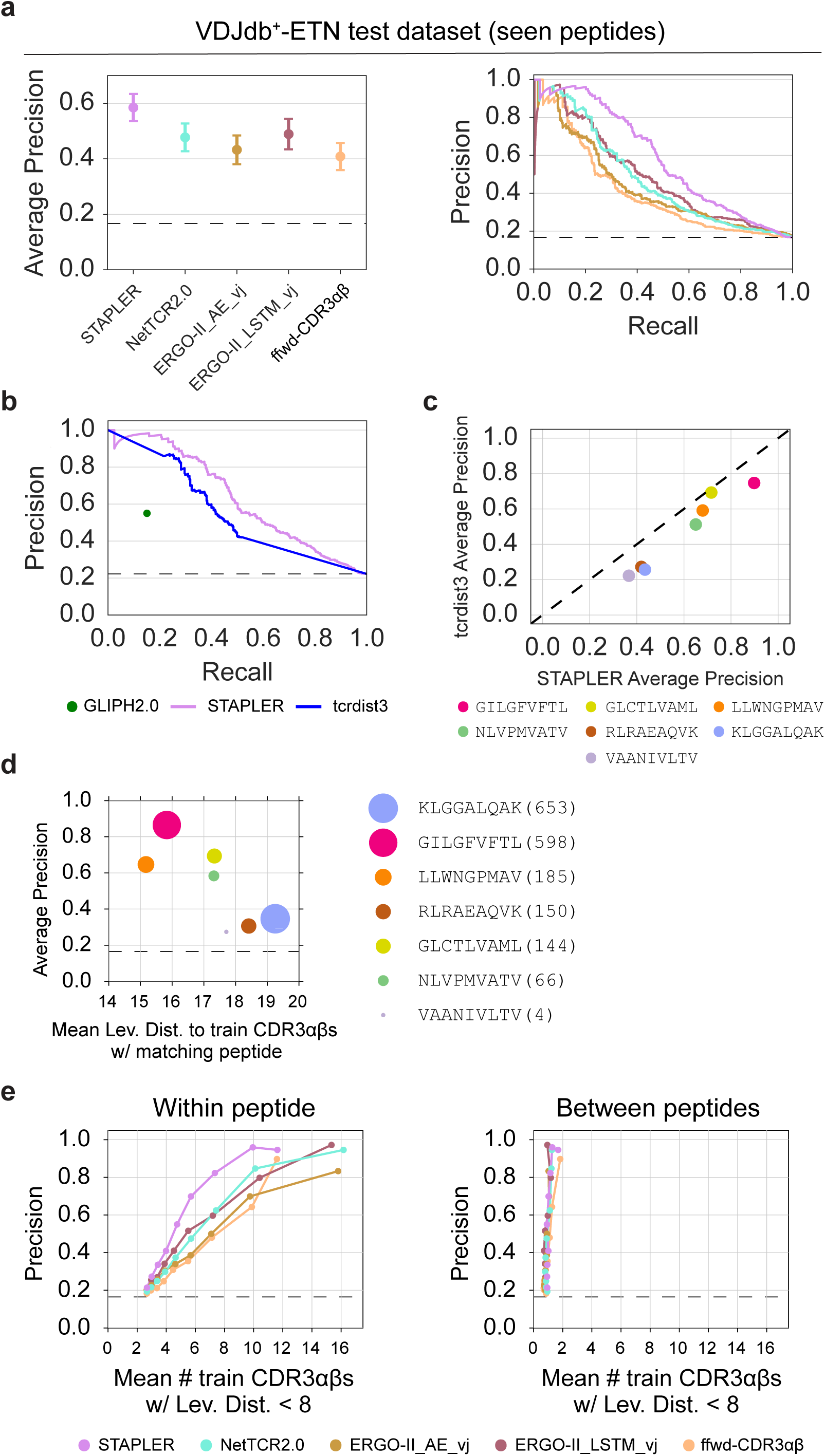
Superior learning efficiency of STAPLER. **a**. Benchmarking of STAPLER against joint TCRαβ-peptide models. Mean average precision (mAP) (left), and precision across recall thresholds (right), on predicting the recognition of seen peptide antigens by unseen TCRs in the VDJdb^+^-ETN test dataset that contains negative pairs generated by mispairing with external TCRs identified in oral cavity squamous cell carcinomas^24^ (367 positive and 1,835 negative pairs, 14 unique seen peptides). Dots show mAP and error bars show mAP 95% confidence intervals. **b**. Benchmarking of STAPLER against sequence distance-based models. Precision across recall thresholds in predicting the recognition of seen peptide antigens by unseen TCRs in the VDJdb^+^-ETN-U test dataset (365 positive and 1,278 negative pairs, 14 unique seen peptides). Note that GLIPH2.0 returns a single clustering that does not include all positive test TCRs and can hence only be used to determine precision at one specific recall, in this case 15.1%. **c**. Comparison of AP of STAPLER and tcrdist3 for individual seen peptide antigens in the VDJdb^+^-ETN-U test dataset. **d**, Relationship between AP for each indicated seen peptide antigen in VDJdb^+^-ETN and the mean Levenshtein distance of positive test dataset CDR3αβ sequences that are paired with those peptides to all positive fine-tuning dataset CDR3αβ sequences that are paired with the same peptide. Dot size indicates abundance of positive TCR-peptide pairs for the indicated seen peptides in the fine-tuning dataset, as also depicted by the numbers in the legend. Data are depicted for all seen test peptides with >5 positive test data points, as AP estimates becomes unstable with fewer data points. **e**. Relationship between precision and the mean number of positive fine-tuning dataset CDR3αβ sequences within a Levenshtein distance of 8 from positive test dataset CDR3αβ sequences in the VDJdb^+^-ETN dataset that are paired with the same seen peptide antigens (left), or, as negative control, are paired with a different peptide (right). See Extended Data Fig. 4b for results when using other Levenshtein distances. To reduce noise in mean Levenshtein distance estimates, precision and mean Levenshtein distance were estimated in increasing steps of 0.1 recall. **c-d**. Levenshtein distance was estimated using the Levenshtein python package^51^. **a**, **b**, **d**, **e**. Dashed line shows random performance (AP of 0.1667).

In prior work, TCRs reactive with known antigens have also been identified with methods that cluster TCRs based on sequence distance metrics, using known TCRs as reference points to assign reactivities. GLIPH2.0^25^ and tcrdist3^26^, two widely used methods to cluster TCRs based on the similarity of their CDR3β (GLIHP2.0) or CDR1αβ, CDR2αβ, CDR2.5αβ, and CDR3αβ (tcrdist3) sequences, have, for instance, been used to identify TCRs reactive to viral antigens. To compare the performance of STAPLER to these distance-based ML models, test dataset TCRs were clustered with seen peptide annotated training dataset TCRs (see Methods). As tcrdist3 computes a TCRαβ distance matrix, the k nearest neighbor algorithm can be used to calculate reactivity scores for each test TCR-peptide pair. In contrast, GLIPH2.0 returns a single clustering solution, and precision can hence only be estimated at the recall of this clustering solution (Methods). GLIPH2.0 was able to annotate the correct peptide label to 15.1% of the positive test TCRs in the VDJdb^+^-ETN-U test dataset that was used for this evaluation (see Methods). At this recall of 15.1%, GLIPH2.0 showed a precision of 55%, while on the same dataset and at the same recall, the precision of STAPLER was 98.2% (**Fig. 3b**). Moreover, STAPLER also outperformed tcrdist3 by +15% AP, and this superior performance was consistent across all tested peptides (**Fig. 3b, 3c**).

As shown above (**Fig. 3a**), STAPLER outperforms neural network and distance-based models in identifying novel TCRs that are reactive with known antigens, suggesting that STAPLER is more efficient at the identification of patterns in limited labeled αβ TCR-peptide data. To evaluate this hypothesis, we first analyzed the relationship between model performance for different epitopes and either the number of TCR-peptide pairs for each epitope that was present during fine-tuning, or the mean similarity of fine-tuning dataset CDR3αβ pairs to test dataset CDR3αβ pairs for each epitope. As shown in **Fig. 3d** and **Extended Data Fig. 4a**, the quality of predictions across epitopes is captured well by the mean similarity of the fine-tuning dataset and test dataset CDR3αβ pairs, providing a rationale to determine the relationship between this mean CDR3αβ pair similarity and precision for the different models. Notably, using this metric, STAPLER was shown to require substantially fewer train TCRs within a given Levenshtein distance to test TCRs to achieve the same prediction performance (**Fig. 3e**), and this more efficient learning was consistent across Levenshtein distances (**Extended Data Fig. 4b**). These data demonstrate that STAPLER displays a superior capacity to learn from limited labeled αβ TCR-peptide data.

## Discussion

The creation of effective TCR – pMHC predictors forms one of the Holy Grails in immunology and depends on two developments: 1. the generation of technology for high-throughput measurement of TCR – pMHC interactions; and 2. modeling approaches that can efficiently learn from the data obtained through these technologies. Here, we present STAPLER: a transformer language model that shows best-in-class performance in prediction of TCR – pMHC reactivity.

STAPLER was developed with the two main aims. First, the incorporation of an MLM pre-training task in model creation allowed the use of the large sets of unlabeled CDR3αβ pairs and MHC-associated peptide sequences that are available. Second, amino acid masking during pre-training and fine-tuning was incorporated with the aim to create a more comprehensive view of interactions within and between TCRαβ and peptide sequences. Interestingly, ablation experiments demonstrated that the superior performance of STAPLER is primarily due to amino acid masking during fine-tuning, in which labeled TCR – pMHC pairs are shown. This contrasts with the value of MLM in natural language processing fields, in which the MLM task is primarily used to increase performance as a self-supervised pre-training task^17^. We offer two potential explanations for this unexpected dependency of STAPLER on amino acid masking during fine-tuning. First, while pre-training was performed using random CDR3αβ-peptide combinations (see *Model pre-training* in the Methods for rationale), full-length TCR sequences were used during fine-tuning. As pMHC recognition is not restricted to CDR3αβ sequences but is non-redundantly encoded in the V and J regions of the full-length TCR^2^, the presence of full length TCR sequences in the MLM fine-tuning task may have allowed the learning of additional (long range) interactions that could not be learned in pre-training. Second, and more speculatively, in the natural language field in which language models were developed, the texts that are offered during pre-training lack a label but are internally coherent. In contrast, in the pre-training phase of STAPLER, the CDR3αβ and the peptide sequence are both internally coherent, but the random combinations of CDR3αβ sequences and peptide sequences are artificially generated. As a result, an MLM pre-training task may learn language representations within CDR3αβ sequences and within peptide sequences, but not between CDR3αβ sequences and peptide sequences. In contrast, as labeled reactive and non reactive TCR-peptide pairs are shown during fine-tuning, the learning of recognition patterns between TCR and peptide sequences by amino acid masking could be feasible at that stage.

A number of wet-lab efforts may be expected to enhance the predictive potential of STAPLER. First, the data used by us and others for model training has been obtained using a number of different methodologies (MHC multimer staining, functional assays etc.), and may not be of consistent quality.

In this regard it is noteworthy that a sizable percentage of TCRs (7.7% of all CDR3αβ data points and 2.7% of all unique CDR3αβ sequences) in the currently available data is labelled as reactive with more than one peptide sequence. While TCRs have the capacity to productively recognize a large number of different antigens^4^, the cross-reactivity as documented within the finite peptide space covered in the current fine-tuning dataset appears unexpected. To improve the quality of the dataset used for model training, validation of all TCRs for which epitopes have currently been mapped using a single standardized screening method could therefore be attractive. Second, to predict the recognition of unknown antigens by unknown TCRs, TCR – pMHC reactivity models will require training on data that covers a substantially larger peptide space. Towards this goal, it may be attractive to consider the dense screening of TCR reactivity in a subset of epitope space that is adjacent to known reactivities. By fine-tuning STAPLER and other models on such local data, the density of datasets required to predict reactivity against unknown antigens throughout epitope space may potentially be estimated. Importantly, such datasets would also address the limitation of the current datasets that only report reactivity with the peptide sequence that happened to form the disease-associated antigen, while from a structural and modeling point of view, knowledge on any of the many other peptide sequences that can be recognized is of equal value. In addition, we note that models generated with such dense local data, may enable the therapeutic engineering of TCRs, for instance with respect to reactivity against self-antigens or, in case of recurring oncogenic mutations, with respect to the capacity to recognize structurally related neoantigens, such as the KRAS codon 12 mutations. As a final goal for the coming years, while TCRs show major differences in the pMHC density required for productive signaling, the annotation of reactivity in the current datasets is binary (i.e., yes, no). It may thus be speculated that datasets that provide a quantitative measure of TCR reactivity will provide a richer source of information for model training.

Given the performance of STAPLER in predicting TCR recognition of known antigens, in particular for many of the viral antigens (e.g., CMV, EBV, influenza A, SARS-Cov2, and Yellow Fever virus), we propose that STAPLER will be of value to identify and characterize virus-specific T cells in the rapidly expanding scRNA-seq datasets of human disease-associated tissues. In line with this, ongoing work suggests the ability to accurately call virus-specific T cell infiltrates in human cancer samples. To allow the use of STAPLER for these and other purposes, the STAPLER model, and code that can be used to fine-tune new (disease-specific) STAPLER models, have been made available at https://github.com/NKI-AI/STAPLER.

## Methods

### Datasets

#### Pre-training dataset

To collect CDR3αβ data for pre-training, TCR sequences were obtained from the following datasets: Yost et al.^27^, Kourtis et al.^28^, Liu et al.^29^, Zheng et al.^30^ and Wu et al.^31^ (**Extended Data Fig. 1b**). Datasets were parsed by extracting CDR3αβ sequences, V and J gene annotations, and CD8 or CD4 expression information. From these datasets, 159,859 pre-train CDR3αβ pairs were selected for which CDR3αβ sequences and V and J gene annotations could be extracted, and CD8 expression was confirmed. To collect 9-mer peptide data for pre-training, 183,398 human MHC class I-associated 9-mer peptides were extracted from the dataset used to train the MHC class-I binding prediction algorithm Anthem^32^.

#### Combined dataset

To collect positive (i.e., reactive) full-length αβ TCR-peptide sequence pairs, we parsed the IEDB^33^, Francis^34^, 10X^35^, McPas^36^, and VDJdb^37^ datasets into a single “combined dataset” (**Extended Data Fig. 1c**). From these datasets, columns that contain the recognized peptide sequence, CDR3α and CDR3β sequences, V and J gene annotations, MHC allele type, and association with either CD8 or CD4 T cells were extracted. For each dataset, only TCR-peptide pairs that contained human TCRs were considered. All duplicate data points within a dataset, containing duplicate entries in each extracted column (including peptide sequence), were removed, retaining only the first datapoint. After combining the different datasets, duplicate data points between datasets were also removed.

VDJdb^37^ is a manually curated and regularly updated database that contains positive TCR-peptide pairs from mouse and human origin. We cloned the VDJdb database from the VDJdb-web GitHub page on 11-10-2021. 10X data present in VDJdb was removed from the database, as the 10X dataset was parsed separately (see below).

The 10X dataset^35^ contains positive αβ TCR-peptide pairs identified by staining 150,000 CD8^+^ T-cells that were isolated from four donors with 44 DNA-barcoded-pMHC-multimers and subsequent sequencing of αβ TCR and DNA-barcodes using 10X V(D)J + Feature-Barcode sequencing. We downloaded the contig annotation and binarized matrix for each of the four donors on 20-10-2021 from the Application Note - A New Way of Exploring Immunity (v1 Chemistry) dataset found at https://www.10xgenomics.com/resources/datasets. Entries were filtered in the contig annotations matrix on high confidence (‘is_cell’) and productive annotation (quality assignments further described in the 10X application note). To create the final 10X dataset, cell barcodes were matched between the contig annotation and binarized matrix, and the separate datasets from the four donors were concatenated.

The McPas database^36^ contains disease-associated positive TCR-peptide pairs of human and mouse origin. We downloaded the McPas dataset from the Friedman lab website on 28-09-2021. Data points that contained remarks on datapoint quality were removed.

The immune epitope database (IEDB)^33^ is a manually curated and regularly updated database that contains, amongst others, positive TCR-peptide pairs from human and non-human origin that are associated with allergy, infectious diseases, transplantation, cancer, and autoimmunity. We downloaded the IEDB dataset on 15-11-2021 using their web-based query function with the following settings: Host: Homo sapiens, linear epitope, MHC-I, and T cells only.

The Francis dataset^34^ contains positive αβ TCR-peptide pairs identified by staining 96,909,416 CD8^+^ T-cells from 27 acute phase and 28 convalescent SARS-CoV-2-infected patients and 23 unexposed individuals with 434 SARS-CoV-2-derived DNA-barcoded-pMHC-multimers, and subsequent sequencing of αβ TCR and DNA-barcodes using 10X V(D)J + Feature-Barcode sequencing. We downloaded the Francis dataset from their supplementary data file S3 (sciimmunol.abk3070_data_file_s3.xlsx) on 18-11-2021. Data coupled to the 188 cell barcodes associated with multiple α chain sequences (5.48% of total) were removed.

#### Filtering of the combined CDR3αβ dataset

After parsing, the combined dataset was filtered to only contain data points with the following properties:

- Human CDR3αβ sequences that start with cysteine (`C`) and end in either phenylalanine (`F`) or tryptophan (`W`), as defined by IMGT^38^. Supplementary Table **2** shows the percentage of CDR3 αβ sequences that was removed because of these `C` and `F/W` quality constraints. Of note, only 9.21% of the CDR3β sequences from IEDB started with a `C`. However, 44.8% of the CDR3β sequences from IEDB ended with a `F` or `W`, and 87.21% of those started with an `A`, suggesting that these CDR3β sequences had been annotated without a starting `C`. We therefore added back the starting `C` for those CDR3β sequences, thereby making the distribution of the starting motifs of these CDR3β sequences similar to those in the other datasets (see Supplementary Table **3** for the starting motifs of CDR3β sequences that end with `F` or `W` in the IEDB dataset before and after correction, and in the other datasets).
- CDR3αβ sequences that contain standard amino acids as defined by IUPAC standards. The percentage of CDR3αβ sequences that was removed because of this constraint is shown in Supplementary Table **2**. From 12 peptides a non-IUPAC suffix (e.g., "LLFGFPVYV + SCM(F5)") was removed.
- CDR3αβ sequences that are paired with 9-mer peptides that bind MHC-I.
- CDR3αβ sequences of which both CDR3 regions have a length between 5-25 amino acids. Note that almost all CDR3αβ sequences fall within this range (**Extended Data Fig. 1d**).

#### Reducing imbalance of the number of CDR3αβ sequences that are paired with each unique peptide in the combined dataset

The number of CDR3αβ sequences that is paired with each unique peptide in this combined dataset is highly imbalanced (**Extended Data Fig. 1e**). For example, CDR3αβ sequences associated with the `KLGGALQAK` (KLG) peptide make up 15,589 of the 22,419 positive TCR-peptide pairs in the combined dataset, while CDR3αβ sequences associated with 342 of the other peptide sequences occur only once. To reduce this imbalance, the following reduction strategy was used (**Extended Data Fig. 1e**). First, MMseqs2 cluster^39^ was used to cluster CDR3αβ sequences that were paired with each unique peptide with the following parameters: `--max-seqs 10000 -c 0.8 --cov-mode 0 -- spaced-kmer-mode 1 --alignment-mode 3 -e 0.001 --min-seq-id 0.9 -- remove-tmp-files 1 -s 4.0 -v 1`. Next, to reduce imbalance, MMseqs2 result2repseq was used to retain only one representative CDR3αβ sequence per cluster. For CDR3αβ sequences paired with all epitopes other than the KLG peptide, CDR3αβ sequences were clustered together when sequences were at least 90% similar (`--min-seq-id 0.9`). For CDR3αβ sequences paired with the KLG peptide, a 54% similarity threshold was used for clustering (`--min-seq-id 0.54`). This more stringent reduction was used to lower the fraction of CDR3αβs paired with the KLG peptide such that negative pair generation by mispairing was feasible for this TCR-peptide combination (see section *Negative data generation by mispairing*).

#### Splitting the combined dataset into a fine-tuning and VDJdb+ test dataset

The combined dataset, as described above, was split into a fine-tuning and a VDJdb^+^ test dataset by moving all positive TCR-peptide pairs from the VDJdb database into a separate dataset (**Extended Data Fig. 1c**). To increase the number of unseen peptides in this VDJdb^+^ test dataset, all positive TCR-peptide pairs of three peptides were included in the VDJdb^+^ test dataset (18 `LLFGYPVYV`, 23 `ALWEIQQVV`, and 48 `SPRWYFYYL` pairs). In addition, to evaluate model performance in predicting the recognition of KLG, the most abundant peptide in the fine-tuning dataset, 95 KLG positive TCR-peptide pairs were included in the VDJdb^+^ test dataset. Finally, 50 positive TCR-peptide pairs of the `RLRAEAQVK` peptide were included, as an example of a peptide that, next to KLG, binds HLA-A*03:01, resulting in a total of 579 positive TCR-peptide pairs (**Extended Data Fig. 1c**).

#### Negative data generation by mispairing

As the resulting fine-tuning and VDJdb^+^ test datasets only contain positive pairs, synthetic negative pairs were separately generated for the fine-tuning and VDJdb^+^ test datasets by mispairing. To generate negative TCR-peptide pairs, peptides from each positive TCR-peptide datapoint were mispaired with five TCRs that were randomly sampled from the set of all TCRs in that dataset that were not originally paired with that peptide, resulting in a total of 19,620 fine-tuning and 2,895 VDJdb^+^ negative TCR-peptide pairs (generating a total of 23,544 fine-tuning and 3,474 VDJdb^+^ TCR-peptide pairs). While TCRs are known to be cross-reactive against millions of different pMHC complexes^4^, in view of the overall size of the pMHC space^5^, the probability of generating false negatives by such mispairing (i.e., generating a TCR-peptide pair that is labeled as negative but would be reactive) is low. Finally, because the combined dataset contained a number of TCRs that were paired with multiple peptides in the individual datasets, mispairing resulted in duplicate αβ TCR-peptide data points between and within the fine-tuning dataset and VDJdb^+^ test dataset. To prevent data leakage from the fine-tuning dataset to VDJdb^+^ test dataset, such duplicate data points (6 positive and 28 negative data points) were removed from the test dataset. As a side note, for the fine-tuning dataset, we also explored the consequence of using external sets of TCRs, rather than internal TCRs, to generate negative pairs by mispairing. In line with prior data^15^, the resulting model showed better-than-random 5-fold cross-validation performance on the fine-tuning dataset when excluding the peptide sequence in the joint TCR-peptide pair input. These data indicate that when using such external TCRs during fine-tuning, resulting models can identify externally sampled TCR sequences as being negative independent from the peptide that these TCR are artificially paired with, and hence the use of negative data points created by mispairing of internal TCRs is preferred for fine-tuning.

To evaluate the effect of data leakage by mispairing with TCRs that were originally paired with seen peptides, we mispaired unseen peptides of each datapoint five times with TCRs sampled from the measured negative TCRs in the DFCI dataset. In addition, seen peptides of each datapoint were mispaired five times with TCRs sampled from an external dataset of 5,021 αβ TCR sequences identified in oral cavity squamous cell carcinomas^24^. This dataset, rather than the DFCI measured negative TCRs, was subsequently used to create negatives for seen peptide data points, because the number of unique measured negative TCRs in DFCI is too low relative to the number of seen peptide data points in the VDJdb^+^ dataset to allow mispairing with the more abundant seen peptides in the VDJdb^+^ database.

#### DFCI external test dataset

To generate the DFCI dataset, all data points that were associated with 9-mer peptides and of which the full-length TCRαβ could be reconstructed were retained. Measured negatives were reconstructed by pairing negative TCRs (326 unique TCRs that did not recognize any peptide in the co-culture experiments) with peptides from the set of unique peptides that was screened in the co-cultures. Reconstruction was performed by pairing the peptide of each of the 132 positive TCR-peptide data points with five of these measured negative TCRs.

#### Reconstruction of full-length αβ TCR sequences in the fine-tuning and test datasets

Full-length αβ TCR sequences excluding the TCRαβ constant domains were reconstructed from CDR3αβ and V and J gene annotations as described in **Extended Data Fig. 1f**. V and J gene annotations were used to look-up IMGT/GENE-DB (IMGT) reference amino acid sequences^38^. In cases in which no V gene allelic annotation was provided, we assumed that the first V gene allele in IMGT was used. Data points without complete V and J gene annotations were removed. To reconstruct full-length αβ TCRs, V gene amino acid sequences were trimmed from their COOH-terminus to the final `C` residue (`C104`). J gene amino acid sequences were trimmed from their NH2-terminus to the first amino acid of the J gene end motif (either `FGXG`, `WGXG`, `XGXG`, or `FXXG`)^38^. Next, full-length αβ TCR sequences were reconstructed by concatenating the trimmed V gene, CDR3, and trimmed J gene sequences. Following reconstruction, a test was performed to determine whether the amino acid and nucleotide V and J IMGT gene annotations were identical. In cases that these annotations differed, the annotation was removed, resulting in the unintentional removal of TRAV IMGT gene annotations for 3 of the 3,924 positive train data points (0.076% of all positive fine-tuning data points). As ERGO-II models, but not STAPLER, uses such V gene annotations, this resulted in a marginally smaller fine-tuning dataset for the ERGO models.

To determine the quality of our TCR reconstruction algorithm, it was evaluated using an internal dataset of 930 manually reconstructed TCR sequences. In addition, we tested the algorithm on a 10X Genomics dataset of 9,299 measured TCRs sequences that were expressed by T cells isolated from peripheral blood mononuclear cells (PBMC)^40^. Sequences reconstructed by the algorithm matched the manually reconstructed sequences in >99% of cases and matched TCR sequences measured by 10X in 84.74% of cases. The differences between the reconstructed and measured sequences were explained by absence of V and J gene allele annotations, potential mutations or sequencing errors, and differences in the reference used for mapping (10X CellRanger uses the Ensembl reference genome for mapping, which differs slightly from the IMGT V- and J amino acids reference sequences).

#### Definition of unseen peptide and unseen TCR sequences

Because peptide diversity is limited in the available labeled αβ TCR-peptide data, we defined unseen peptides in the test datasets as peptides whose sequences do not exactly match any peptide sequence in the fine-tuning dataset. At present, there is no consensus on what constitutes an unseen TCR when that TCR is paired with a seen peptide^41^. We defined unseen positive test TCRs that were paired to a seen peptide in the VDJdb^+^ dataset as having CDR3αβ sequences that were less than 90% similar (54% for the KLG peptide) to any positive fine-tuning dataset CDR3αβ sequence that was paired to the same seen peptide, as estimated by MMseqs2^39^ Smith-Waterman alignment. We defined TCRs from the external TCR dataset and TCRs that are or were (before negative generation by mispairing) paired to unseen peptides in DFCI and VDJdb^+^ dataset as “unseen” when their CDR3αβ sequences did not exactly match any sequence in the fine-tuning dataset.

### Development of STAPLER

STAPLER is inspired by the BERT model^17^ that uses the encoder from the transformer model^16^, and learns deep bidirectional language representations using the MLM task. We implemented the encoder from transformers model using the x-transformers package^42^. From the original encoder we changed the ReLU activation function to a GLU activation in every feedforward sub-layer of the encoder^43^ and we set the dropout at 5% for all dropout layers (after the learned embedding, after each feedforward and self-attention layer in the encoder, and in the CLS pooling layer).

#### Input representation

Input amino acid sequences were tokenized using a vocabulary of 20 amino acids and 5 special tokens (classification token, separation token, masking token, padding token, unknown token). A classification token ([CLS]) that is used as an aggregate TCR – peptide pair sequence representation for predicting whether that pair is positive or negative (i.e., the prediction of recognition label task) was included as the first token of the input. A separation token ([SEP]) was included between the TCRα and peptide sequence and between the peptide and TCRβ chain sequence. In every epoch, 15% of randomly chosen input amino acid tokens was replaced by a masking token ([MASK]). Masking was used for both the pre-training and fine-tuning MLM tasks, no masking was used during validation/testing. A padding token ([PAD]) was used to pad inputs to match the length of the longest TCR-peptide pair sequence. Finally, any character in the TCRα, TCRβ, or peptide sequences that is not formed by one of the naturally occurring amino acids is replaced by the unknown token ([UNK]). Note that in the analyses described here, any sequences containing such characters were removed from the fine-tuning and test datasets, and the UNK token was hence not used. Each token is encoded as a 25-dimensional vector, initialized randomly. During fine-tuning this embedding was optimized (i.e., a learnable embedding), changing the embedding values of each token based on relationships between tokens. Additionally, because a transformer is invariant with respect to the reordering of the input^44^ we added the T5 relative positional encoding^45^.

#### Model pre-training

Pre-training was, for two reasons, performed on CDR3αβ sequences rather than full-length TCRs. First, pre-training using MLM on unlabeled data is only useful when it allows the use of a larger dataset in pre-training than the labeled data available for fine-tuning, which is the case for the large publicly available dataset of paired CDR3α – CDR3β sequences. Second, sequence diversity is not homogeneously divided over TCR sequences (i.e., with CDR3 regions showing a substantially higher diversity than V and J gene regions), making it challenging to simultaneously perform an MLM task with a constant masking rate on both CDR3 and V/ J gene regions.

Model pre-training was performed in 500 epochs using the MLM task. In each iteration in these epochs, a new collection of 159,859 random CDR3αβ-peptide combinations were shown. These random CDR3αβ-peptide combinations were made by pairing each of the 159,859 collected pre training CDR3αβ sequences to a peptide randomly sampled from the 183,398 collected pre-training peptides that bind MHC-I. To calculate the loss for the MLM task, the final representation of each MASK token was transformed to an output vector with a dimension of 25, which corresponds to the size of the token vocabulary (see Input representation). The cross-entropy loss was then calculated between this predicted probability distribution over all tokens in the vocabulary and the true distribution, which is a one-hot vector with a value of 1 for the original token and 0 for all other amino acids in the vocabulary.

#### Model fine-tuning

After pre-training, the parameters of the pre-trained model were fine-tuned using 5-fold cross validation on the 23,544 labeled fine-tuning pairs of CD8^+^ T cell-derived full-length αβ TCRs and 9-mer peptides that bind MHC-I, using both the prediction of recognition label task and the MLM task (**Fig. 1c**). We trained STAPLER for 150 epochs and selected the best STAPLER model based on AP on the validation dataset (see *Evaluation on the test datasets*).

The MLM task during fine-tuning was identical to the MLM task used in pre-training. The prediction of recognition label task involved the transformation of the final representation of the CLS token to an output vector with a dimension of two (i.e., positive and negative), and the subsequent calculation of the cross-entropy loss between the output vector and the recognition label, which is either 1.0 (i.e., “positive”) or 0 (i.e., “negative”) for a given TCR-peptide pair. The loss for a single input sequence was defined as the sum of the CLS (ℒ_cls_) and MLM losses (ℒ_mlm_) that are scaled by their corresponding, optimized (see below), scaling factors.

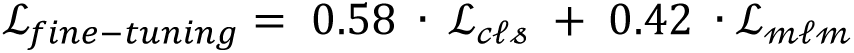

During fine-tuning, positive TCR-peptide pairs were oversampled, and negative TCR-peptide pairs were undersampled, such that the ratio of positive to negative TCR-peptide pairs was equal. Furthermore, to oversample peptide sequences that were paired with few TCRs and undersample peptide sequences that were paired with many TCRs (e.g., KLG), unique peptide sequences were sampled with a probability that is relative to a sampling weight calculated as the log2 ratio of the inverse number of positive and negative TCR-peptide pairs of each unique peptide sequence (equal to the log2 of the ratio between the number of data points of the fine-tuning dataset and the number of data points for that unique peptide sequence). To determine the effectiveness of this weighted random sampling approach, the number of positive and negative TCR-peptide pairs that were shown during fine-tuning were visualized (data not shown). In addition, we visualized the number of pairs shown for each unique peptide sequence during fine-tuning. This visualization demonstrated that the number of positive and negative pairs shown during fine-tuning was equal, and that the imbalance in the number of positive and negative TCR-peptide pairs per unique peptide sequence shown during fine-tuning was reduced. It is noted that when peptide sequence imbalance is not reduced during fine tuning, performance of the model in 5-fold cross-validation shows a further increase (data not shown), an observation that may be explained by a superior performance on those peptide sequences that are abundant in both the fine-tuning and validation folds. Thus, in applications of STAPLER in which superior performance of the model in predicting recognition of the most abundant peptides is of interest, omission of the weighted sampling approach should be considered. For validation and testing, no weighted random sampling was used.

For both pre-training and fine-tuning, a batch size of 256 and learning rate of 0.005 was used. The Adam optimizer that is used in the original BERT paper^17^ was replaced with the AdamW optimizer^46^.

In addition, the learning rate scheduler was replaced from a linear warmup with linear decay to a linear warmup (10 epochs) with cosine decay.

##### Optimization

First, performance of three model sizes, Tiny, Mini, and Medium (with the previously described specifications^17^) was compared (**Extended Data Fig. 2b**). The number of heads, layers and hidden dimension of the multi-head attention encoder layer determines the number of parameters of the model^16^, and model parameters can be found in Supplementary Table **4**. The Medium model showed a slightly improved performance over the other models, and this model size was therefore selected for STAPLER.

To determine the optimal hyperparameters for fine-tuning, the dropout, learning rate, mask percentage, and scaling factors of the MLM task and prediction of recognition label task losses were optimized for the ‘no pre-training Mini sized’ and the ‘no pre-training & fine-tuning CDR3αβ Mini sized’ models (**Extended Data Fig. 2b**). Hyperparameter optimization was performed using 100 separate 5- fold cross-validation runs, and hyperparameters were optimized for the highest mAP over the 5-folds using the Bayesian search algorithm Tree-structured Parzen Estimator (TPE)^47^ (implemented in the Optuna^48^ hyperparameter tuning package). Early stopping was implemented with a patience of 30 epochs (stopping fine-tuning when the AP on the validation fold did not increase after 30 epochs).

The optimized ‘no pre-training Mini sized’ model outperformed the optimized ‘no pre-training & fine tuning CDR3αβ Mini sized’ model, and considering optimization being resource intensive, the optimal hyperparameters of this model were used for fine-tuning of the pre-trained Medium sized model (i.e., STAPLER). The selected hyperparameters are described in Supplementary Table **4**.

### Baseline ffwd, ERGO-II, and NetTCR-2.0 models

To create a baseline model, we trained a ffwd model on the CDR3αβ-peptide sequences of the 23,544 labeled fine-tuning pairs. Input representation was similar to that described for STAPLER, except that tokens were not masked, as a ffwd model cannot be trained using the MLM task. The length of the first layer of the ffwd was set to the maximal length of the CDR3ab-peptide input sequences (i.e., 60 tokens). To match input layer length, all input sequences shorter than this maximum length were padded to a length of 60 tokens. After matching the input layer length, we flattened the input and passed it through a series of 5 fully connected layers. The first 4 layers had an input and output size of 294, each followed by a ReLu activation. The final layer had an input size of 294 and output size of 2, followed by a ReLu activation. Additionally, a 15% dropout layer was applied between the fourth and final layer. We calculated the cross-entropy loss between the final 2-dimensional output vector and the recognition label, which is either 1.0 (i.e., “positive”) or 0 (i.e., “negative”) for a given TCR-peptide pair. A batch size of 4,096 with a learning rate of 0.003 was used, and layers, hidden dimension, dropout % and learning rate were optimized similar to what is described for STAPLER. The total number of parameters of the ffwd model was 0.75M.

The ERGO-II^12^ directory and the NetTCR2.0^11^ directory were cloned from GitHub on 12-07-2022 and 22-02-2022. Both models were trained using their default parameters.

### Model performance evaluation

#### 5-fold cross-validation on the fine-tuning dataset

The performance of STAPLER, NetTCR-2.0, ERGO-II, and the baseline ffwd models was estimated using AP in 5-fold cross-validation on the fine-tuning dataset. For each of the 5 iterations, all models were trained on exactly the same folds and validated on the same held-out validation fold. AP was estimated for each of the 5 validation folds using the average_precision_score function from Scikit-learn^49^. Reported values are the median AP of the 5 validation folds.

#### Evaluation on the test datasets

After training five versions of each of the STAPLER (fine-tuned), ERGO-AE-vj, ERGO-LSTM-vj, and ffwd models using 5-fold cross-validation, the final prediction score for each of these models was calculated on the test datasets (e.g., VDJdb^+^ and DFCI) by taking the average of the prediction scores returned by each of these five versions. NetTCR-2.0 was trained five times on the complete fine-tuning dataset, and the final prediction score was calculated as the average of these five versions. This adjustment was required as NetTCR-2.0 always selects the model that was trained for 100 epochs over the whole fine-tuning dataset, while all other models select the best model version using highest AP on the validation fold during 5-fold cross-validation. In addition, as NetTCR-2.0 uses different random seeds for each training run, training NetTCR-2.0 five times was also required to exclude that performance improvements were solely due to a particular random seed. The mean and the 95% confidence intervals of mAP on the test sets were estimated by 1,000-fold bootstrapping. It is noted that the widely used AUROC metric will be disproportionally inflated in cases of data leakage when such leakage leads to an increased ability to predict the more abundant negative pairs (i.e., the majority class), as AUROC is sensitive to class imbalance^50^ (**Extended Data Fig. 3f-h**).

### Comparison of STAPLER, GLIPH2.0, and tcrdist3 in predicting TCR recognition of known antigens

Performance of STAPLER, GLIPH2.0^25^, and tcrdist3^26^ in predicting the recognition of seen (known) peptides in the VDJdb^+^-ETN test dataset was compared as follows. To annotate GLIPH2.0 and tcrdist3 clusters with known pMHC specificities, clustering of test dataset TCRs that recognize seen peptides was performed together with the TCRs from positive pairs in the fine-tuning dataset. First, duplicate TCRαβs were removed from the VDJdb^+^-ETN dataset, generating the VDJdb^+^-ETN-U test dataset (leaving 365 positive and 1,278 negative unique test dataset TCRs) and fine-tuning dataset (leaving 3,621 positive fine-tuning TCRs). Next, TCRs that were shared between the resulting sets of fine-tuning and test TCRαβs were removed from the fine-tuning dataset (removing 5 TCRs).

The resulting sets of unique fine-tuning and test dataset TCRαβs (n=5,259 TCRs) were combined to generate the final input for GLIPH2.0 clustering. Using default settings (CD8 reference version 1.0), the GLIPH2.0 clustering website tool (accessed at http://50.255.35.37:8080 on 16/03/2023) was able to cluster 16% of the positive and negative unique test dataset TCRs (392 test TCRs). All test dataset TCRs that GLIPH2.0 assigned to clusters that did not contain any fine-tuning dataset TCRs (159 test TCRs; this also includes ‘single’ pattern clusters) were removed, as annotation of TCRs requires the presence of a fine-tuning dataset TCR in the same cluster. Note that the removal of positive test dataset TCRs by this step and other steps below reduces recall but favors precision, as this is calculated within the set of test dataset TCRs that is predicted to recognize a given peptide (i.e., predicted to be “positive”). To subsequently annotate test dataset TCRs that were clustered together with fine-tuning dataset TCRs, we used majority voting of the peptides that are recognized by those fine-tuning dataset TCRs (voting being required in 39.9% of clusters, in the remaining clusters all fine-tuning dataset TCRs were annotated with the same peptide). Majority voting could not be used for 49 test dataset TCRs present in clusters that had a tied peptide vote, and these TCRs were removed (note that the random picking of a peptide for these TCRs would have reduced the precision estimate of GLIPH2.0). TCRs can be clustered into multiple clusters by GLIPH2.0 and, because of this, 25 test dataset TCRs were annotated with different peptides. As we observed that the separate counting of each of these peptide annotations decreased the precision estimate of GLIPH2.0, these 25 test TCRs were also removed. Of the remaining 159 annotated test dataset TCRs, 46 test dataset TCRs were removed that were annotated with a peptide that is not present in the VDJdb^+^-ETN-U test dataset. Finally, to allow for comparison to STAPLER and tcrdist3, 13 positive test dataset TCRs were removed that were annotated with another peptide than the peptide that these TCRs were paired with in the VDJdb^+^-ETN-U test dataset (thereby also increasing the precision estimate of GLIPH2). This resulted in 100 predicted test dataset TCRs of which 55 were true positives and 45 false positives, leading to a precision of 55%, at a recall of 15.1% (55 true positives identified amongst a total of 365 test dataset positives).

We used tcrdist3 to compute distances between TCRαβ sequence pairs in both the resulting sets of unique fine-tuning and test dataset TCRαβs. Fine-tuning dataset TCRαβ distances were then used to train a KNN classifier with 3 nearest neighbors using the KNeighborsClassifier function from Scikit learn^49^. The predict_proba method of the trained KNeighborsClassifier was then used to predict a score for each test dataset TCRαβs, using the computed test dataset TCRαβ distances as input. This score represents the probability of the reactivity of those test dataset TCRαβs to their paired peptide, with higher scores indicating a greater likelihood of reactivity.

To compare the performance of GLIPH2 and tcrdist3 to that of STAPLER, we also used STAPLER to predict reactivity scores for the 1,643 TCRαβ-peptide pairs in VDJdb+-ETN-U dataset that were used to evaluate GLIPH2 and tcrdist3. As both STAPLER and tcrdist3 output a continuous reactivity score for each predicted TCR-peptide pair, precision can be plotted as a function of recall at different score thresholds. In contrast, GLIPH2.0 returns a binary prediction (a positive test TCR can be annotated or not by the default clustering solution) and precision can therefore only be calculated at a single specific recall (in this case 15.1%).

Table S1: Previously established TCR-pMHC specificity models

Table S2: Quality of CDR3αβ sequences in datasets before selection

Table S3: Adding a starting C to the CDR3β sequences in IEDB

Table S4: Model sizes and parameters

## Supporting information

Table S1 - Previously established TCR-pMHC specificity models

Table S2 - Quality of CDR3ab sequences in datasets before selection

Table S3 - Adding a starting C to the CDR3b sequences in IEDB

Table S4 - Model sizes and parameters

## Acknowledgements

This work was supported by the Stevin award and Louis Jeantet prize, and by the Institute for Chemical Immunology (to T.N.S.) and by an institutional grant of the Dutch Cancer Society and the Dutch Ministry of Health, Welfare and Sport. We would like to thank Joris van de Haar and Benoit Nicolet for their feedback on the manuscript.

## Author Contributions

B.P.Y.K., M.M., and T.N.S. conceived and designed the study. T.N.S. and M.M. co-supervised the study.

B.P.Y.K. and M.M. prepared the final the training and test datasets. B.P.Y.K. implemented and developed the STAPLER model in consultation with M.M. B.P.Y.K. and M.M. evaluated the STAPLER model and generated the figures. G.O. and C.J.W. provided data and interpreted results. J.T. and E.M provided computational resources, advised on practical model implementation, model training, model optimization, and model evaluation, and interpreted results. M.M., B.P.Y.K., and T.N.S. wrote the manuscript. All authors reviewed and revised the manuscript.

---

**Extended Data Fig. 1:**
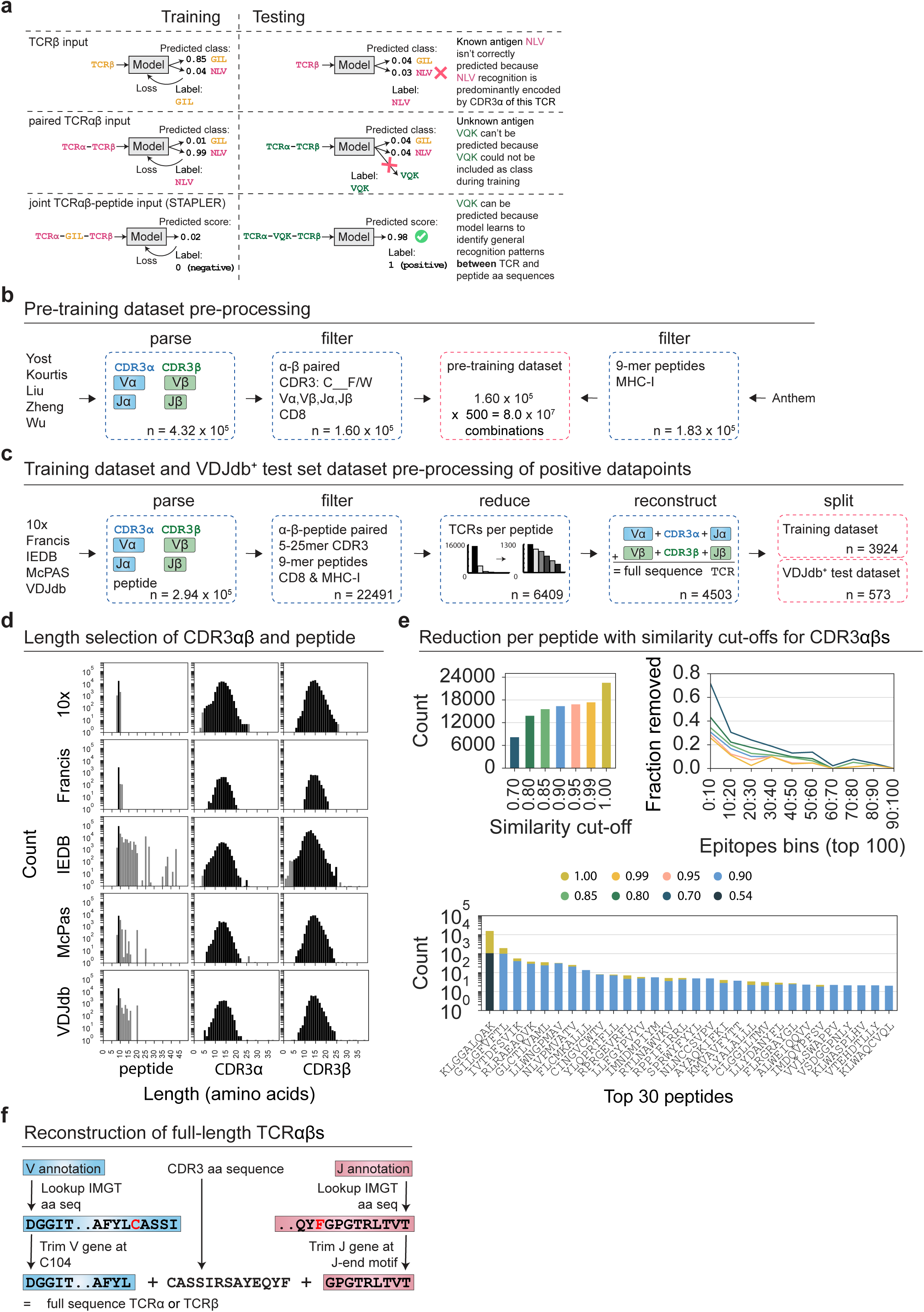
Datasets. **a.** A joint TCRαβ-peptide input is preferable for predicting the recognition of both known (i.e., present in the fine-tuning dataset) and unknown antigens, as (top) pMHC recognition is non-redundantly encoded in both the TCRα and β chain^2^, and (top and middle) models that solely use the TCR as input can only be trained to predict reactivity against predetermined peptide antigen categories. As argued previously^10–15^, models with a joint TCRαβ and peptide input (bottom) have the potential to predict recognition of both known and unknown categories. The value of such joint encoding has likewise been shown in vision models that jointly encode text labels and images to predict whether these are matched, thereby not being restricted to a predetermined set of image categories^52^. **b**. CDR3αβ sequences for pre-training were acquired by: (i) parsing αβ TCR sequences from the indicated single-cell TCR αβ sequencing datasets, and (ii) selecting paired CDR3αβ sequences that start with a `C` and end in either a `F` or `W`, have V and J gene annotations, and annotated CD8 expression. 9-mer peptides for pre-training were acquired from the dataset used to train the Anthem prediction model^32^. **c.** The labeled full length αβ TCR-peptide fine-tuning dataset and VDJdb^+^ test dataset were acquired by: (i) parsing the indicated datasets/databases that store positive TCR-peptide pairs; (ii) selecting data points that contain paired CDR3αβ sequences that have: a length between 5 and 25 amino acids (both CDR3 regions, see **d**), V and J gene annotation, annotated CD8 expression, and are paired with 9-mer peptides; (iii) reduction of the imbalance of CDR3αβ sequences that were paired with each unique peptide sequence (Methods); (IV) reconstruction of full length αβ TCR sequences (Methods); and (V) splitting of the combined dataset into a fine-tuning and a VDJdb^+^ test dataset (Methods). **d**. Length distribution of selected (colored) and excluded (grey) peptide (left), CDR3α (middle), and CDR3β (right) sequences in the indicated datasets/databases. **e**. CDR3αβ sequences that were paired with each unique peptide sequence were reduced using maximal similarity thresholds of either 99%, 95%, 90%, 85%, 80% or 70%. The total number of CDR3αβ sequences left after reduction when using either threshold (top left) and the fraction of CDR3αβ sequences removed from the set of CDR3αβ sequences paired with each peptide (top right) are depicted. Data are depicted for the top 100 peptides, binned in groups of 10 by the number of CDR3αβ sequences that these peptides were paired with. A threshold of 90% was chosen for all peptides other than `KLGGALQAK` for which a similarity threshold of 54% was used in order to reduce TCR number sufficiently to allow for negative data generation by mispairing. The total number of CDR3αβ sequences paired with the top 30 most abundant peptides in the dataset after reduction with different similarity thresholds is depicted (bottom). **f**. Reconstruction of full length αβ TCR sequences using their V and J annotations and CDR3αβ amino acid sequences. V and J gene segment annotations were used to look-up amino acid reference sequences in the IMGT/GENE-DB database. V segment amino acid reference sequences were trimmed from their COOH-terminus to the final `C` residue (`C104`), and J segment amino acid reference sequences were trimmed from their NH_2_-terminus to the first amino acid of the J gene end motif^38^. Finally, full-length TCR chains were reconstructed by concatenating trimmed V, CDR3, and trimmed J sequences.

**Extended Data Fig. 2:**
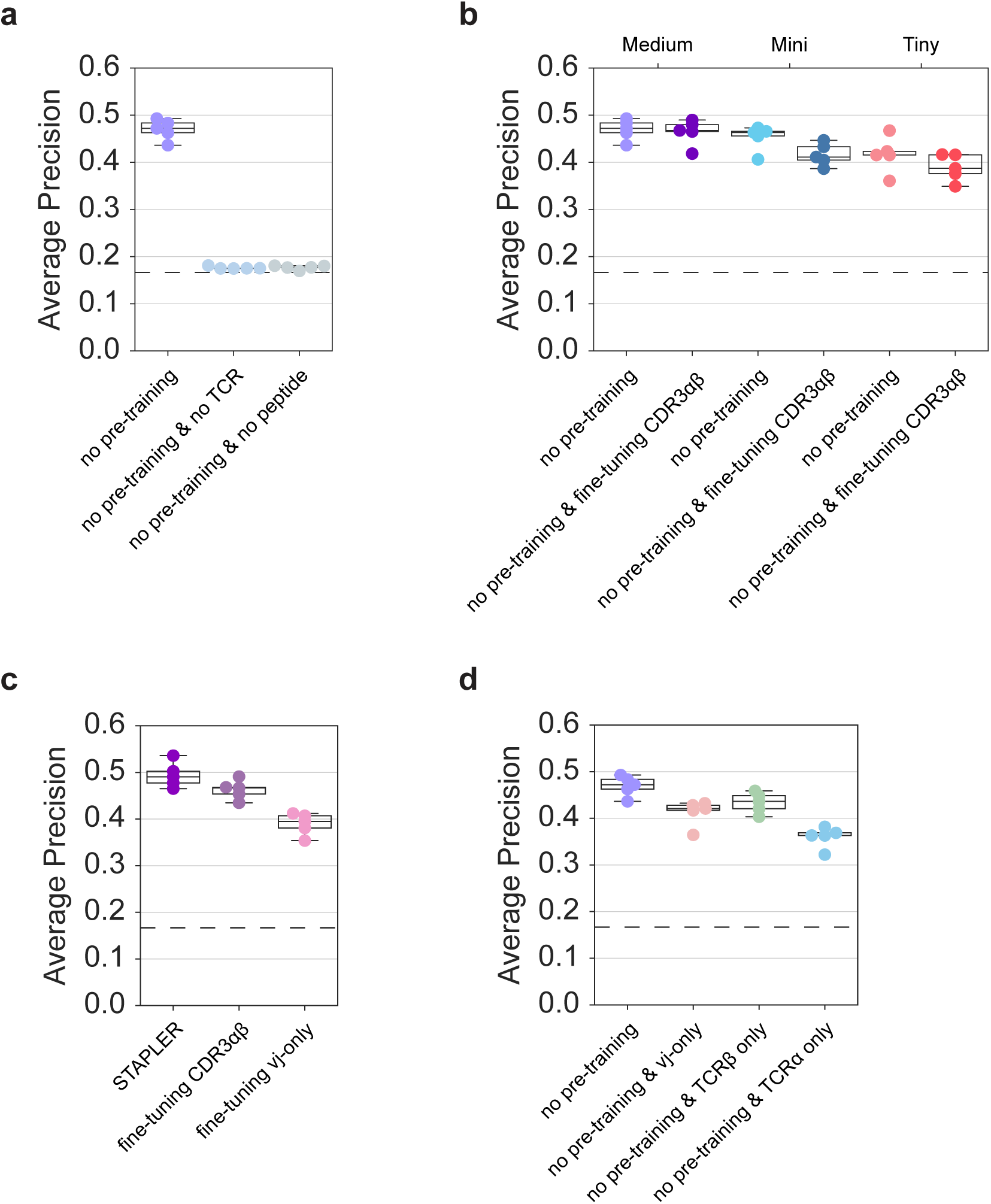
Validation of STAPLER using 5-fold cross-validation experiments. **a**. Average precision (AP) of indicated model variants of STAPLER in 5-fold cross-validation on the fine-tuning dataset. Models were compared to variants of STAPLER from which either the TCRαβ sequences (to control for bias in peptide sequences) or peptide sequences (to control for bias in TCR sequences) were removed from the input. For simplicity, all models compared here were created without pre-training component. **b**. Comparison of AP of indicated model variants of STAPLER that have different standardized BERT model sizes^17^, and either using full-length TCRαβ sequences or CDR3αβ sequences as input for fine-tuning. For simplicity, all models compared here were created without pre-training component. **c**. Comparison of mean AP of STAPLER and STAPLER models that were either fine-tuned on only CDR3αβ sequences, or on full-length αβ TCR sequences without CDR3αβ sequences (V and J sequence only). **d**. Comparison of AP of indicated model variants of STAPLER that were either fine-tuned on full-length αβ TCR sequences, full-length αβ TCR sequences without CDR3αβ sequence (V and J sequence only; “no pre-training & vj-only”), β chain sequences (“no pre-training & TCRβ only”), or α chain sequences (“no pre-training & TCRα only”). For simplicity, all models compared here were created without pre-training component. Note that both TCRα and TCRβ sequences contribute to model performance. **a-d**. Dots show AP of each fold, boxes indicate model AP median and 25th/75th percentiles, whiskers show min/max except for outliers determined using the inter quartile range (IQR). Dashed line shows random performance (AP of 0.1667).

**Extended Data Fig. 3:**
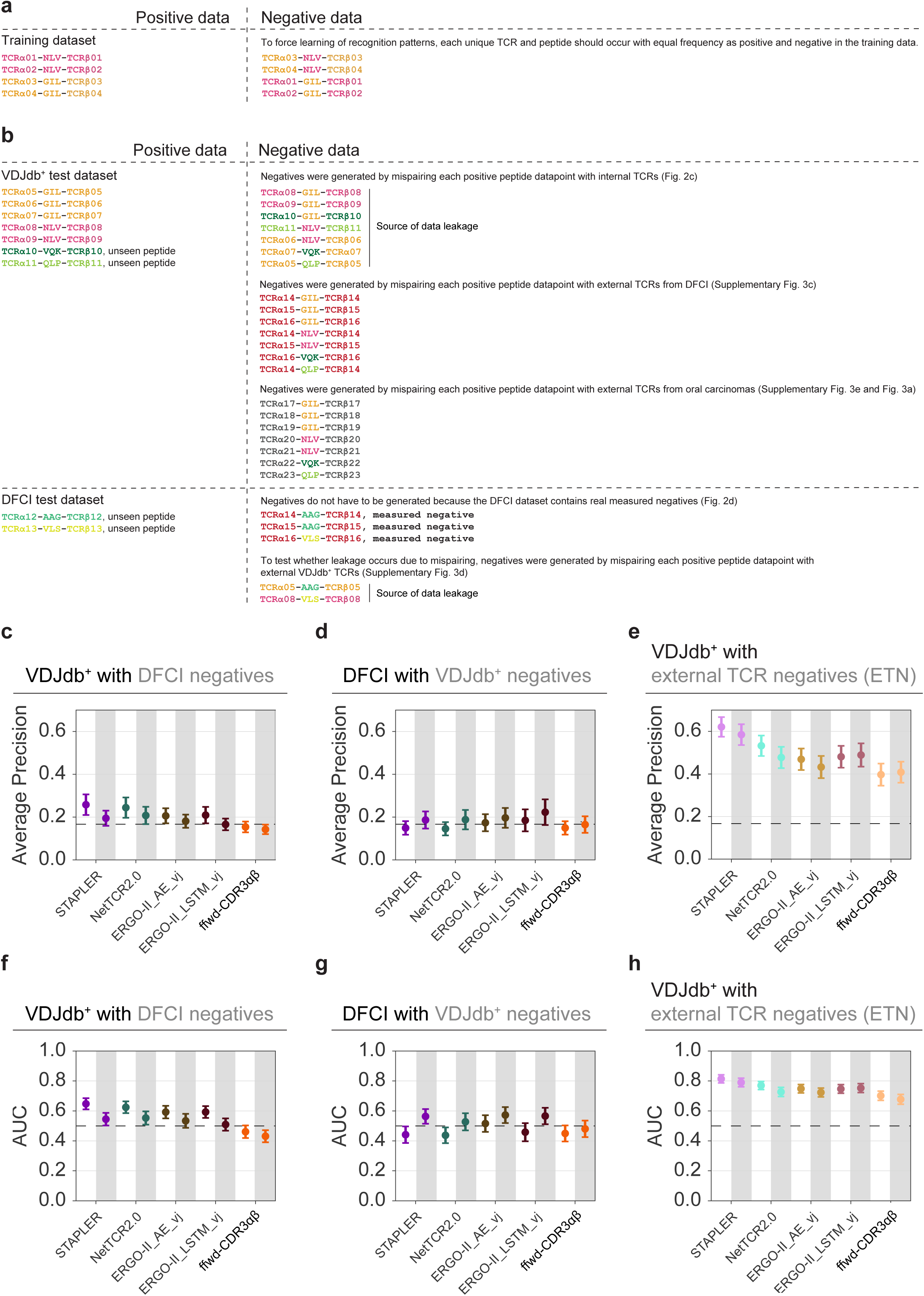
Data leakage as a confounder in performance evaluation of TCR – pMHC specificity models. **a**. Avoidance of class occurrence bias. To force learning of recognition patterns, rather than preferential occurrence in either the negative or positive class, each unique TCR and each unique peptide sequence occurs with equal frequency as positive and negative in the training data. **b**. Employed strategies for negative data generation. In all depicted mispairing strategies, TCRs were sampled from the set of all TCRs that were not paired to same peptide sequence as the positive peptide datapoint that was being mispaired. Also, in all depicted mispairing strategies, the sampling probability of TCRs was equal to their abundance in the dataset from which those TCRs were sampled. Figure numbers shown between brackets indicate evaluation of model performance on datasets that contain negative data generated by the mispairing strategy that is depicted. Note that strategies in which test dataset negatives are generated by mispairing each positive peptide datapoint with TCRs that are related to TCRs that are abundant in the fine-tuning dataset (see Fig. 2c, Extended Data Fig. 3d) results in an overestimate of model performance, due to class L3.2 data leakage^20^. **c**. Mean average precision (mAP) on predicting recognition of unseen peptide antigens by unseen TCRs in the VDJdb^+^ test dataset with negatives either being generated by mispairing peptides of each positive unseen peptide datapoint with internal TCRs from its positive pairs (white), or by mispairing each positive unseen peptide datapoint with TCRs from the set of negative TCRs from the measured negative pairs in the DFCI dataset (grey shading). **d**. mAP on predicting recognition of unseen peptide antigens by unseen TCRs in the DFCI dataset containing either measured DFCI negatives (white), or with negatives being generated by mispairing each positive unseen peptide datapoint with TCRs from the positive pairs in VDJdb^+^ (grey shading). **e**. mAP on predicting recognition of seen peptide antigens by unseen TCRs in the VDJdb^+^ test dataset with negatives either being generated by mispairing each positive seen peptide datapoint with internal TCRs from its positive pairs (white), or by mispairing each positive unseen peptide datapoint with TCRs that were identified in oral cavity squamous cell carcinomas^24^ (grey shading). **c-e**. Dots show mAP and error bars show the mAP 95% confidence interval. Dashed line shows random performance (0.1667 AP). **f**. Area Under the ROC curve (AUC) on predicting recognition of unseen peptide antigens by unseen TCRs in the VDJdb^+^ test dataset with negatives either being generated by mispairing each positive unseen peptide datapoint with internal TCRs from its positive pairs (white), or with TCRs from the set of negative TCRs from the measured negative pairs in the DFCI dataset (grey shading). **g**. AUC on predicting recognition of unseen peptide antigens by unseen TCRs in the DFCI dataset containing either measured DFCI negatives (white), or with negatives being generated by mispairing each positive unseen peptide datapoint with TCRs from the positive pairs in VDJdb^+^ (grey shading). **h**, AUC on predicting recognition of seen peptide antigens by unseen TCRs in the VDJdb^+^ test dataset with negatives either being generated by mispairing each positive seen peptide datapoint with internal TCRs from its positive pairs (white), or with TCRs that were identified in oral cavity squamous cell carcinomas^24^ (grey shading). **f-h**. Dots show mean AUC and error bars show the AUC 95% confidence interval. Dashed line in shows random performance (0.50 AUC). Negative data generation strategies for (**c-e**) and (**f**-**h)** are illustrated in (**b**).

**Extended Data Fig. 4:**
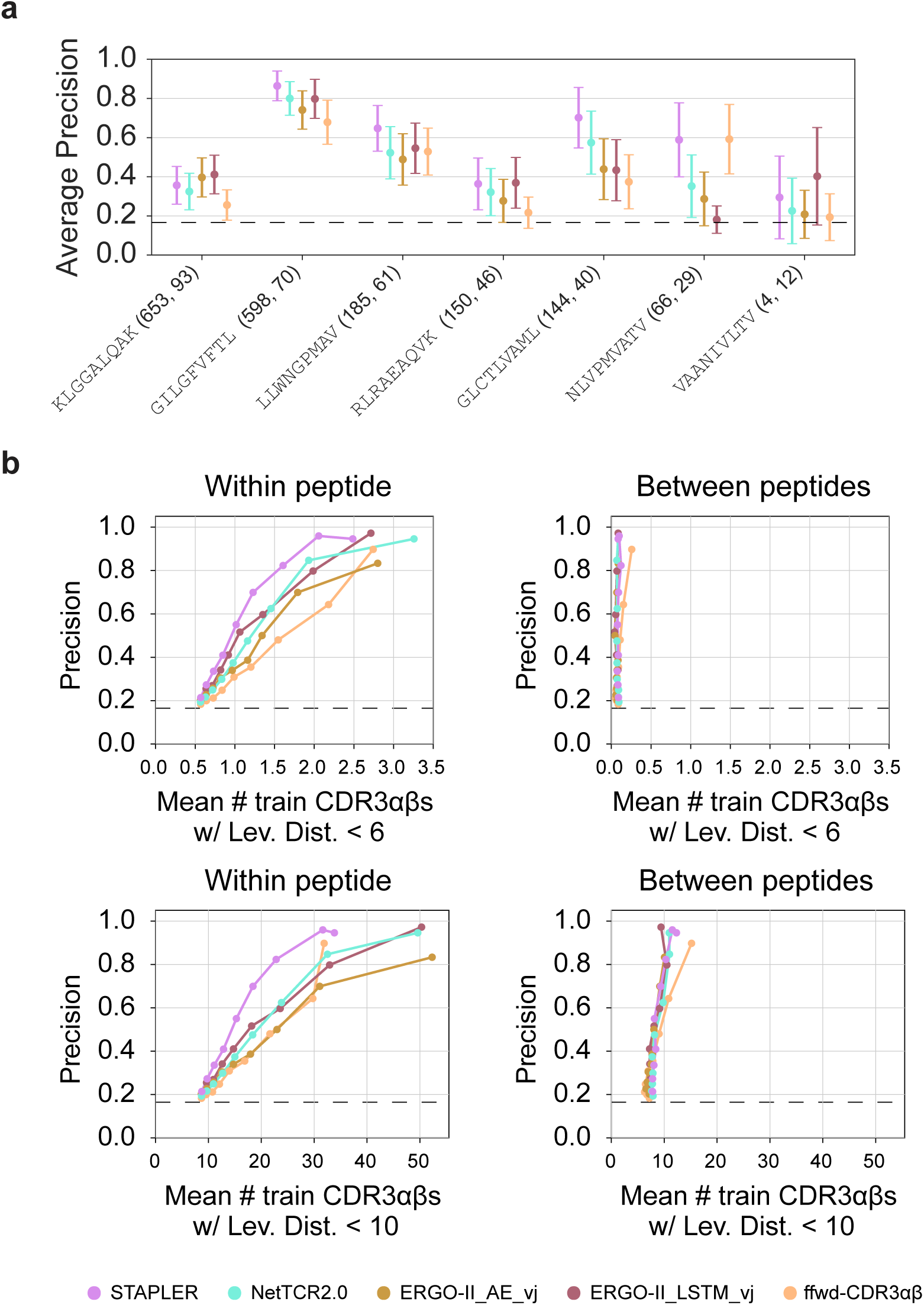
Predictive performance and benchmarking of STAPLER. **a**. Mean average precision (mAP) for seen peptides in the VDJdb^+^-ETN test dataset that contains negatives generated by mispairing with external TCRs identified in oral cavity squamous cell carcinomas^24^. Numbers after peptide sequences indicate the number of positive data points for that peptide in the fine-tuning dataset, and the number of positive data points for that peptide in the VDJdb^+^-ETN test dataset. Data are depicted for all seen test peptides with >5 positive test data points, as AP estimates becomes unstable with fewer data points. Dots show mAP and error bars show the mAP 95% confidence interval. **b**. Relationship between precision and the mean number of positive fine-tuning dataset CDR3αβ sequences within a Levenshtein distance of either 6 (top) or 10 (bottom) from positive test CDR3αβ sequences that are paired with seen peptide antigens that either match (left), or, as negative control, do not match (right) the peptides of the positive fine-tuning dataset CDR3αβ sequences. To reduce noise in mean Levenshtein distance estimation, precision and mean Levenshtein distance were estimated in increasing steps of 0.1 recall. Levenshtein distance was estimated using the Levenshtein python package^51^. **a**-**b** Dashed line shows random performance (AP of 0.1667).

## References

1. Gaud, G., Lesourne, R. & Love, P. E. Regulatory mechanisms in T cell receptor signalling. Nat Rev Immunol 18, 485–497 (2018).

2. Milighetti, M., Shawe-Taylor, J. & Chain, B. Predicting T Cell Receptor Antigen Specificity From Structural Features Derived From Homology Models of Receptor-Peptide-Major Histocompatibility Complexes. Front Physiol 12, 1358 (2021).

3. Mora, T. & Walczak, A. M. How many different clonotypes do immune repertoires contain? Current Opinion in Systems Biology vol. 18 104–110 Preprint at https://doi.org/10.1016/j.coisb.2019.10.001 (2019).

4. Wooldridge, L. et al. A single autoimmune T cell receptor recognizes more than a million different peptides. Journal of Biological Chemistry 287, 1168–1177 (2012).

5. Sarkizova, S. et al. A large peptidome dataset improves HLA class I epitope prediction across most of the human population. Nat Biotechnol 38, 199–209 (2020).

6. Kaech, S. M. & Cui, W. Transcriptional control of effector and memory CD8+ T cell differentiation. Nature Reviews Immunology 2012 12:11 12, 749–761 (2012).

7. Schumacher, T. N., Scheper, W. & Kvistborg, P. Cancer Neoantigens. Annu Rev Immunol 37, 173–200 (2019).

8. Herold, K. C., Vignali, D. A. A., Cooke, A. & Bluestone, J. A. Type 1 diabetes: translating mechanistic observations into effective clinical outcomes. Nature Reviews Immunology 2013 13:4 13, 243–256 (2013).

9. Le Bert, N. et al. SARS-CoV-2-specific T cell immunity in cases of COVID-19 and SARS, and uninfected controls. Nature 584, 457–462 (2020).

10. Fischer, D. S., Wu, Y., Schubert, B. & Theis, F. J. Predicting antigen specificity of single T cells based on TCR CDR 3 regions . Mol Syst Biol 16, (2020).

11. Montemurro, A. et al. NetTCR-2.0 enables accurate prediction of TCR-peptide binding by using paired TCRα and β sequence data. Commun Biol 4, 1–13 (2021).

12. Springer, I., Tickotsky, N. & Louzoun, Y. Contribution of T Cell Receptor Alpha and Beta CDR3, MHC Typing, V and J Genes to Peptide Binding Prediction. Front Immunol 12, 1436 (2021).

13. Weber, A., Born, J. & Rodriguez Martínez, M. TITAN: T-cell receptor specificity prediction with bimodal attention networks. Bioinformatics 37, i237–i244 (2021).

14. Cai, M., Bang, S., Zhang, P. & Lee, H. ATM-TCR: TCR-Epitope Binding Affinity Prediction Using a Multi-Head Self-Attention Model. Front Immunol 13, (2022).

15. Moris, P. et al. Current challenges for unseen-epitope TCR interaction prediction and a new perspective derived from image classification. Brief Bioinform 22, 1–12 (2021).

16. Vaswani, A. et al. Attention Is All You Need. Adv Neural Inf Process Syst 2017-December, 5999–6009 (2017).

17. Devlin, J., Chang, M. W., Lee, K. & Toutanova, K. BERT: Pre-training of deep bidirectional transformers for language understanding. in NAACL HLT 2019 - 2019 Conference of the North American Chapter of the Association for Computational Linguistics: Human Language Technologies - Proceedings of the Conference vol. 1 4171–4186 (Association for Computational Linguistics (ACL), 2019).

18. Geirhos, R. et al. Shortcut learning in deep neural networks. Nat Mach Intell 2, 665–673 (2020).

19. Dens, C., Bittremieux, W., Affaticati, F., Laukens, K. & Meysman, P. Interpretable deep learning to uncover the molecular binding patterns determining TCR–epitope interactions. bioRxiv 2022.05.02.490264 (2022) doi:10.1101/2022.05.02.490264.

20. Kapoor, S. & Narayanan, A. Leakage and the Reproducibility Crisis in ML-based Science.

21. Chicco, D. Ten quick tips for machine learning in computational biology. BioData Min 10, 1–17 (2017).

22. Oliveira, G. et al. Phenotype, specificity and avidity of antitumour CD8+ T cells in melanoma. Nature 596, 119–125 (2021).

23. Lu, T. et al. Deep learning-based prediction of the T cell receptor–antigen binding specificity. Nature Machine Intelligence 2021 3:10 3, 864–875 (2021).

24. Luoma, A. M. et al. Tissue-resident memory and circulating T cells are early responders to pre-surgical cancer immunotherapy. Cell 185, 2918–2935.e29 (2022).

25. Huang, H., Wang, C., Rubelt, F., Scriba, T. J. & Davis, M. M. Analyzing the Mycobacterium tuberculosis immune response by T-cell receptor clustering with GLIPH2 and genome-wide antigen screening. Nat Biotechnol 38, 1194–1202 (2020).

26. Mayer-Blackwell, K. et al. TCR meta-clonotypes for biomarker discovery with tcrdist3: identification of public, HLA-restricted SARS-CoV-2 associated TCR features. bioRxiv (2021) doi:10.1101/2020.12.24.424260.

27. Yost, K. E. et al. Clonal replacement of tumor-specific T cells following PD-1 blockade. Nature Medicine 2019 25:8 25, 1251–1259 (2019).

28. Kourtis, N. et al. A single-cell map of dynamic chromatin landscapes of immune cells in renal cell carcinoma. Nature Cancer 2022 3:7 3, 885–898 (2022).

29. Liu, T. et al. Single cell profiling of primary and paired metastatic lymph node tumors in breast cancer patients. Nature Communications 2022 13:1 13, 1–17 (2022).

30. Zheng, L. et al. Pan-cancer single-cell landscape of tumor-infiltrating T cells. Science (1979) 374, (2021).

31. Wu, T. D. et al. Peripheral T cell expansion predicts tumour infiltration and clinical response. Nature 579, 274–278 (2020).

32. Mei, S., et al.. Anthem: A user customised tool for fast and accurate prediction of binding between peptides and HLA class i molecules. Brief Bioinform 22, (2021).

33. Vita, R. et al. The Immune Epitope Database (IEDB): 2018 update. Nucleic Acids Res 47, D339– D343 (2019).

34. Francis, J. M. et al. Allelic variation in class I HLA determines CD8 + T cell repertoire shape and cross-reactive memory responses to SARS-CoV-2 MGH COVID-19 Collection and Processing Team. Sci. Immunol vol. 7 https://www.science.org (2022).

35. 10X Genomics. A New Way of Exploring Immunity - Linking Highly Multiplexed Antigen Recognition to Immune Repertoire and Phenotype - 10x Genomics. (2019).

36. Tickotsky, N., Sagiv, T., Prilusky, J., Shifrut, E. & Friedman, N. McPAS-TCR: a manually curated catalogue of pathology-associated T cell receptor sequences. Bioinformatics 33, 2924–2929 (2017).

37. Bagaev, D. V. et al. VDJdb in 2019: Database extension, new analysis infrastructure and a T- cell receptor motif compendium. Nucleic Acids Res 48, D1057–D1062 (2020).

38. Lefranc, M. P. et al. IMGT R, the international ImMunoGeneTics information system R 25 years on. Nucleic Acids Res 43, D413–D422 (2015).

39. Steinegger, M. & Söding, J. MMseqs2 enables sensitive protein sequence searching for the analysis of massive data sets. Nature Biotechnology *2017 35:11* 35, 1026–1028 (2017).

40. 10k Human PBMCs, 5’ v2.0, Chromium X - 10x Genomics. https://www.10xgenomics.com/resources/datasets/10-k-human-pbm-cs-5-v-2-0-chromium-x-2-standard-6-1-0.

41. Meysman, P., et al. Benchmarking solutions to the T-cell receptor epitope prediction problem: IMMREP22 workshop report. bioRxiv 2022.10.27.514020 (2022) doi:10.1101/2022.10.27.514020.

42. Phil Wang. lucidrains/x-transformers: A simple but complete full-attention transformer (release 0.22.1). https://github.com/lucidrains/x-transformers.

43. Google, N. S. GLU Variants Improve Transformer. (2020) doi:10.48550/arxiv.2002.05202.

44. Dufter, P., Schmitt, M. & Schütze, H. Position Information in Transformers: An Overview. Computational Linguistics 48, 733–763 (2021).

45. Raffel, C. et al. Exploring the Limits of Transfer Learning with a Unified Text-to-Text Transformer. Journal of Machine Learning Research 21, 1–67 (2019).

46. Loshchilov, I. & Hutter, F. Decoupled Weight Decay Regularization. 7th International Conference on Learning Representations, ICLR 2019 (2017) doi:10.48550/arxiv.1711.05101.

47. Bergstra, J., Bardenet, R., Bengio, Y. & Kégl, B. Algorithms for Hyper-Parameter Optimization. Adv Neural Inf Process Syst 24, (2011).

48. Akiba, T., Sano, S., Yanase, T., Ohta, T. & Koyama, M. Optuna: A Next-generation Hyperparameter Optimization Framework. Proceedings of the ACM SIGKDD International Conference on Knowledge Discovery and Data Mining 2623–2631 (2019) doi:10.48550/arxiv.1907.10902.

49. Pedregosa, F. et al. Scikit-learn: Machine Learning in Python. Journal of Machine Learning Research 12, 2825–2830 (2011).

50. Saito, T. & Rehmsmeier, M. The Precision-Recall Plot Is More Informative than the ROC Plot When Evaluating Binary Classifiers on Imbalanced Datasets. PLoS One 10, e0118432 (2015).

51. Max Bachmann. Levenshtein . Preprint at https://github.com/maxbachmann/Levenshtein (2021).

52. Radford, A. et al. Learning Transferable Visual Models From Natural Language Supervision.

